# pmultiqc: An open-source, lightweight, and metadata-oriented QC reporting library for MS proteomics

**DOI:** 10.1101/2025.11.02.685980

**Authors:** Qi-Xuan Yue, Chengxin Dai, Selvakumar Kamatchinathan, Chakradhar Bandla, Henry Webel, Asier Larrea, Wout Bittremieux, Julian Uszkoreit, Tom David Müller, Jinqiu Xiao, Juergen Cox, Philip Ewels, Vadim Demichev, Oliver Kohlbacher, Timo Sachsenberg, Chris Bielow, Mingze Bai, Yasset Perez-Riverol

## Abstract

The increasing scale and complexity of proteomics data demand robust, scalable, and interpretable quality control (QC) frameworks to ensure data reliability and reproducibility. Here, we present pmultiqc, an open-source Python package that standardizes and generates web-based QC reports across multiple proteomics data analysis platforms. Built on top of the widely adopted MultiQC framework, pmultiqc offers specialized modules tailored to mass spectrometry workflows, with full initial support for quantms, DIA-NN, MaxQuant/MaxDIA, and mzIdentML/mzML-based pipelines. The package computes a wide range of QC metrics, including raw intensity distributions, identification rates, retention time consistency, and missing value patterns, and presents them in interactive, publication-ready reports. By leveraging sample metadata in the SDRF format, pmultiqc enables metadata-aware QC and introduces, for the first time in proteomics, QC reports and metrics guided by standardized sample metadata. Its modular architecture allows easy extension to new workflows and formats. Alongside comprehensive documentation and examples for running pmultiqc locally or integrated into existing workflows, we offer a cloud-based service that enables users to generate QC reports from their own data or public PRIDE datasets.

## Introduction

Mass spectrometry (MS)-based proteomics experiments are inherently complex, involving multiple stages ranging from sample preparation and instrument acquisition to data processing and statistical analysis. Each of these steps introduces potential sources of technical variability, noise, or systematic bias that can significantly impact the accuracy, reproducibility, and biological interpretability of the data (1). As proteomics increasingly contributes to large-scale biological research and integrative multiomics studies, ensuring the quality of the data has become essential (1, 2). Without rigorous and standardized quality control (QC), the risk of drawing misleading conclusions increases, particularly when integrating proteomics with other omics layers such as transcriptomics or metabolomics, where unreliable data can compromise the entire analysis framework. Consequently, QC is not merely a best practice, but a fundamental requirement for trustworthy proteomic analyses, especially in the context of public repositories deposition and reanalysis (2, 3).

A wide range of tools has been developed to support QC in proteomics (4–7). For instance, PTXQC (6) provides an in-depth evaluation of MaxQuant outputs, rawDiag (8) supports method optimization and diagnostics for raw LC-MS data, PRIDE Inspector Toolsuite (5) provided QC for PRIDE database submissions (3, 5); and MSstatsQC (7) offers a statistical framework for longitudinal monitoring of instrument performance using the MSstats framework. While each of these tools offers unique strengths, they often focus on specific pipelines or stages in the workflow and lack a unified reporting format for the results data or the QC information. In addition, the majority of these tools and packages have been focused on data-dependent acquisition (DDA) workflows, but no major library supports popular data-independent acquisition (DIA) tools such as DIA-NN (9) or MaxDIA (10). Moreover, most existing tools do not natively support standardized sample metadata, such as that provided in SDRF (Sample and Data Relationship Format) (11), which limits their utility in multi-sample or multi-omics contexts where consistent metadata is essential for meaningful comparisons.

To address these limitations, we introduce pmultiqc (pmultiqc.quantms.org), an open-source Python package designed to standardize and generate web-based QC reports across diverse proteomics workflows. Built on top of the widely adopted MultiQC framework (12), pmultiqc takes advantage of its modular and extensible architecture to parse output files from popular proteomics software such as DIA-NN (9), MaxQuant (13) / MaxDIA (10), mzIdentML + mzML (PRIDE Complete submissions), and quantms (14). This integration allows users to visualize and assess key QC metrics, including identification rates, intensity distributions, or retention time consistency, within a single, interactive, and publication-ready report. A distinguishing feature of pmultiqc is its novel use of SDRF-formatted sample metadata as a central component of QC report generation. pmultiqc is an open-source package (https://github.com/bigbio/pmultiqc), also released in PyPI, Bioconda (15), and BioContainers packages (16). In addition to the pmultiqc library, we introduce the pmultiqc service (e.g. https://www.ebi.ac.uk/pride/services/pmultiqc/), to enable research groups to deploy pmultiqc for its users, enabling to generate QC reports from their own data and from public data hosted in the PRIDE database (3). pmultiqc has the potential to become a key resource for transparent, reproducible, and scalable quality assessment in both individual and large-scale collaborative proteomics projects.

### Experimental procedures

#### pmultiqc: core processing flow and data integration

The pmultiqc (https://pmultiqc.quantms.org) processing framework consists of three main stages: (1) data detection and parsing, (2) data integration and QC metric computation, and (3) HTML report generation (**Figure 1A-C**). During the data detection phase, pmultiqc automatically identifies input file types by matching them against registered file format patterns and command-line options, enabling appropriate downstream processing pathways for different workflows and analysis tools (e.g., MaxQuant, **Figure 1A-B**). The parsing stage employs format-specific parsers optimized for memory efficiency and workflow-specific requirements. Each parser extracts only the relevant information needed to generate QC metrics for the corresponding workflow, avoiding unnecessary memory overhead. For example, when processing MaxQuant results, pmultiqc selectively loads peptide information from the *evidence.txt* and *proteinGroups.txt* files rather than loading complete datasets into memory, focusing specifically on data required for MaxQuant QC report generation. This selective parsing approach enables efficient processing of large-scale proteomics datasets while maintaining comprehensive QC coverage. QC metrics (**Figure 1B**) are then transformed into standardized formats compatible with MultiQC’s visualization functions, including bar graphs, line graphs, scatter plots, heatmaps, and interactive tables (**Figure 1C**). The final stage involves sequential data processing through MultiQC’s rendering engine to generate self-contained HTML reports with embedded JavaScript for dynamic visualizations and interactive exploration.

**Figure 1:**
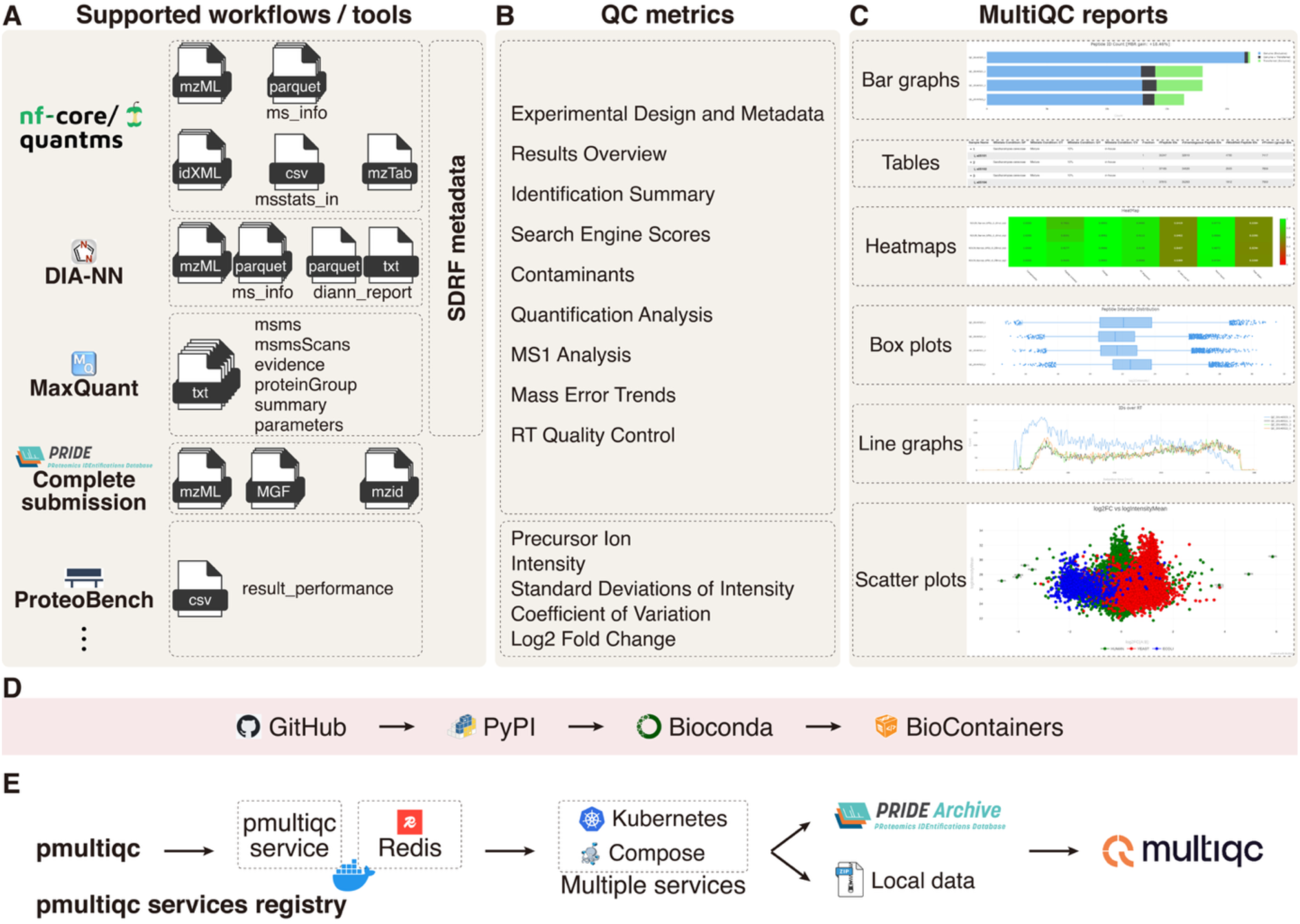
Overview of the pmultiqc proteomics quality control and analysis workflow. (A) The diagram illustrates the integration of multiple proteomics software tools and their output files into a comprehensive quality control pipeline. Supported workflows include quantms, DIA-NN, MaxQuant, ProteomeXchange Complete Submission, and ProteoBench, which generate various output files including mass spectrometry data (*ms_info.parquet, mztab, msstats.csv*), identification files (*id.idxml, diann_report.parquet*), and result summaries (*msms.txt, evidence.txt, proteinGroup.txt, summary.txt, mzidentml, mzML/MGF files,* and *result_performance.csv*). (B) These files are processed through SDRF metadata integration to produce a comprehensive quality control report containing multiple analysis modules: Experimental Design and Metadata, Results Overview, Identification Summary, Search Engine Scores, Contaminants, Quantification Analysis, MS1 Analysis, MS2 and Spectral Stats, Mass Error Trends, RT Quality Control, and various QC metrics including Precursor Ion Intensity, Standard Deviation of Intensity, Coefficient of variation and log2FC calculations. (C) The pmultiqc platform generates diverse visualization outputs, including bar graphs, tables, line graphs, heatmaps, box plots, and scatter plots for comprehensive data interpretation. (D) The pmultiqc library follows FAIR principles through an automated deployment workflow: code is maintained on GitHub, PyPI packages are generated for each release, followed by Bioconda and BioContainer distributions for enhanced accessibility and reproducibility. (E) The pmultiqc service can be deployed in scalable cloud infrastructures using Redis for caching and orchestrated through Docker Compose or Kubernetes, with seamless integration to the PRIDE Archive proteomics database, enabling both local and remote data access for comprehensive proteomics quality assessment.

#### Supported data formats and workflows

pmultiqc supports, by 2025, for four major proteomics analysis workflows and the ProteoBench format, each with specific input file requirements (**Table 1**). For quantms workflows, required files include *experimental_design.tsv* (SDRF-derived metadata), mzTab files (identification and quantification results), MSstats input files (statistical analysis inputs), and *ms_info.parquet* (MS level 1/MS level 2 (MS1/MS2) signal summaries), and optional idXML and YAML parameter files. MaxQuant workflows utilize *parameters.txt* (analysis parameters), *proteinGroups.txt* (protein-level results), *summary.txt* (experiment overview), *evidence.txt* (peptide evidence), and MS/MS scan files (*msms.txt*, *msmsScans.txt*). DIA-NN workflows primarily use *report.tsv* or *report.parquet* files as main inputs, with optional *ms_info.parquet*.

**Table 1:**
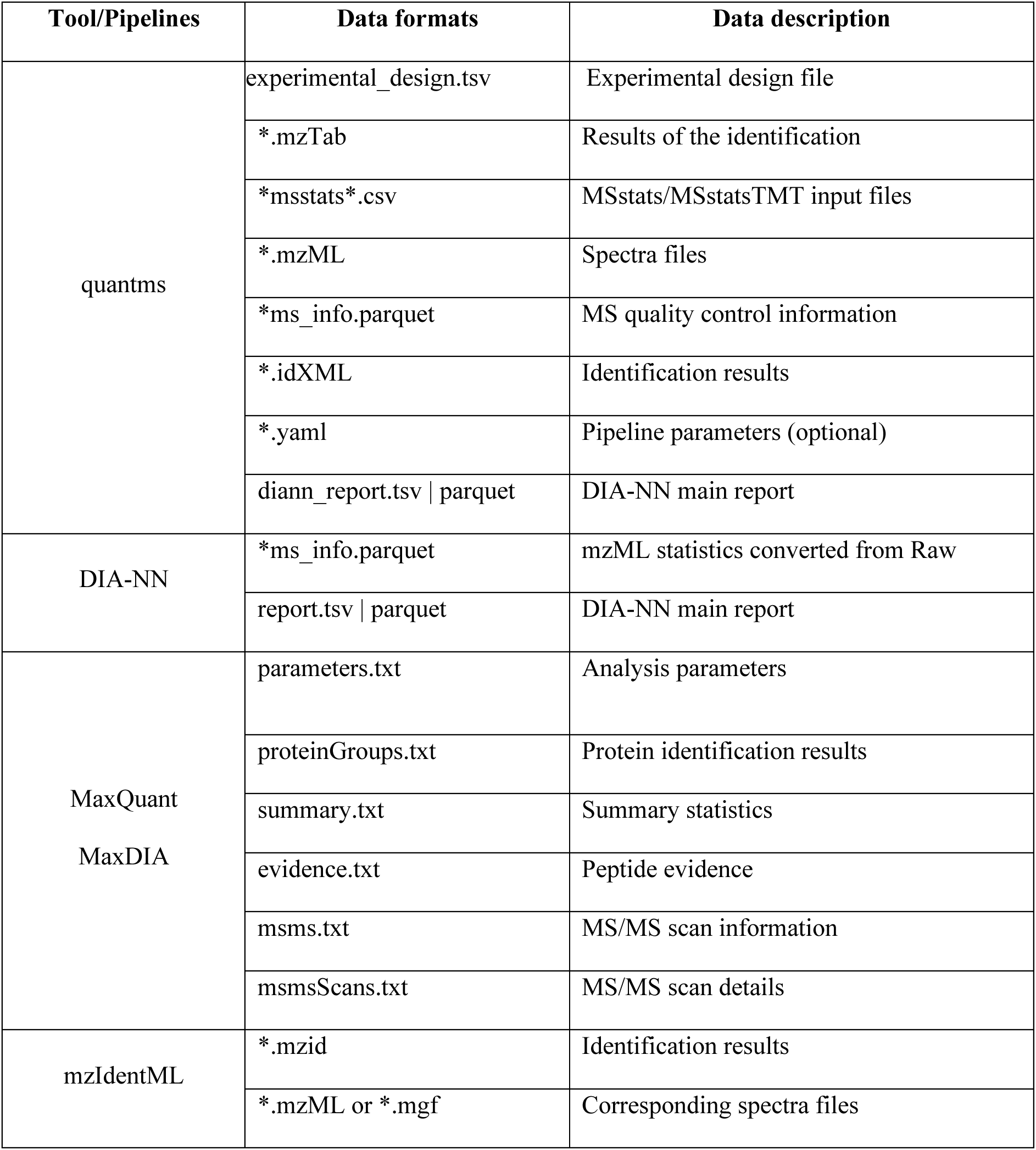
Supported proteomics analysis workflows and pipelines by pmultiqc. The table includes the name of the analysis tool, the output files supported and the description of the information that those files contain. (*) means the name of the file for the given project.

| Tool/Pipelines | Data formats | Data description |
| --- | --- | --- |
| quantms | experimental_design.tsv | Experimental design file |
|  | *.mzTab | Results of the identification |
|  | *msstats*.csv | MSstats/MSstatsTMT input files |
|  | *.mzML | Spectra files |
|  | *ms_info.parquet | MS quality control information |
|  | *.idXML | Identification results |
|  | *.yaml | Pipeline parameters (optional) |
|  | diann_report.tsv \| parquet | DIA-NN main report |
| DIA-NN | *ms_info.parquet | mzML statistics converted from Raw |
|  | report.tsv \| parquet | DIA-NN main report |
| MaxQuant<br>MaxDIA | parameters.txt | Analysis parameters |
|  | proteinGroups.txt | Protein identification results |
|  | summary.txt | Summary statistics |
|  | evidence.txt | Peptide evidence |
|  | msms.txt | MS/MS scan information |
|  | msmsScans.txt | MS/MS scan details |
| mzIdentML | *.mzid | Identification results |
|  | *.mzML or *.mgf | Corresponding spectra files |

#### pmultiqc: library based on MultiQC

pmultiqc is developed as a Python-based extension of MultiQC (https://seqera.io/multiqc/) (12), leveraging its proven extensibility framework for bioinformatics quality control reporting. MultiQC’s plugin architecture enables the development of custom modules without modifying its core codebase, allowing for seamless integration through Python’s entry points mechanism. A plugin/package was developed instead of modules within MultiQC due to pmultiqc’s requirements for proteomics-specific Python dependencies. pmultiqc is automatically discovered and loaded during MultiQC execution, inheriting all the robust features of the MultiQC framework while extending its capabilities specifically for proteomics workflows. pmultiqc is distributed under an open-source license via PyPI (Python Package Index), Bioconda (15), BioContainers (16), and GitHub (https://github.com/bigbio/pmultiqc) (**Figure 1D**). pmultiqc utilizes a variety of specialized libraries for proteomics data processing, including PyOpenMS (17) for handling MS file formats, pyteomics (18) for proteomics-specific data structures, and standard scientific Python libraries including pandas (data manipulation), scikit-learn (statistical analysis), pyarrow (efficient data storage), and NumPy (numerical computations) (19).

## Results

Each bioinformatics workflow and the corresponding pmultiqc reports provide different plots and QC metrics tailored to its input and result data (**Figure 1A-C)**. For example, MaxQuant provides standardized results for identifying contaminants at the peptide, protein, or post-translational modified (PTM) peptide levels (https://pmultiqc.quantms.org/PXD003133/multiqc_report.html#contaminants). We organize the QC reports and metrics into nine main sections similar to PTXQC and PRIDE Inspector Toolsuite reports (see examples in **Table 2**). Each section addresses different aspects of MS data analysis and contains multiple subsections (**Supplementary Notes 1**). It is important to note that the number of sections in the report may vary depending on the specific data used.

**Table 2:**
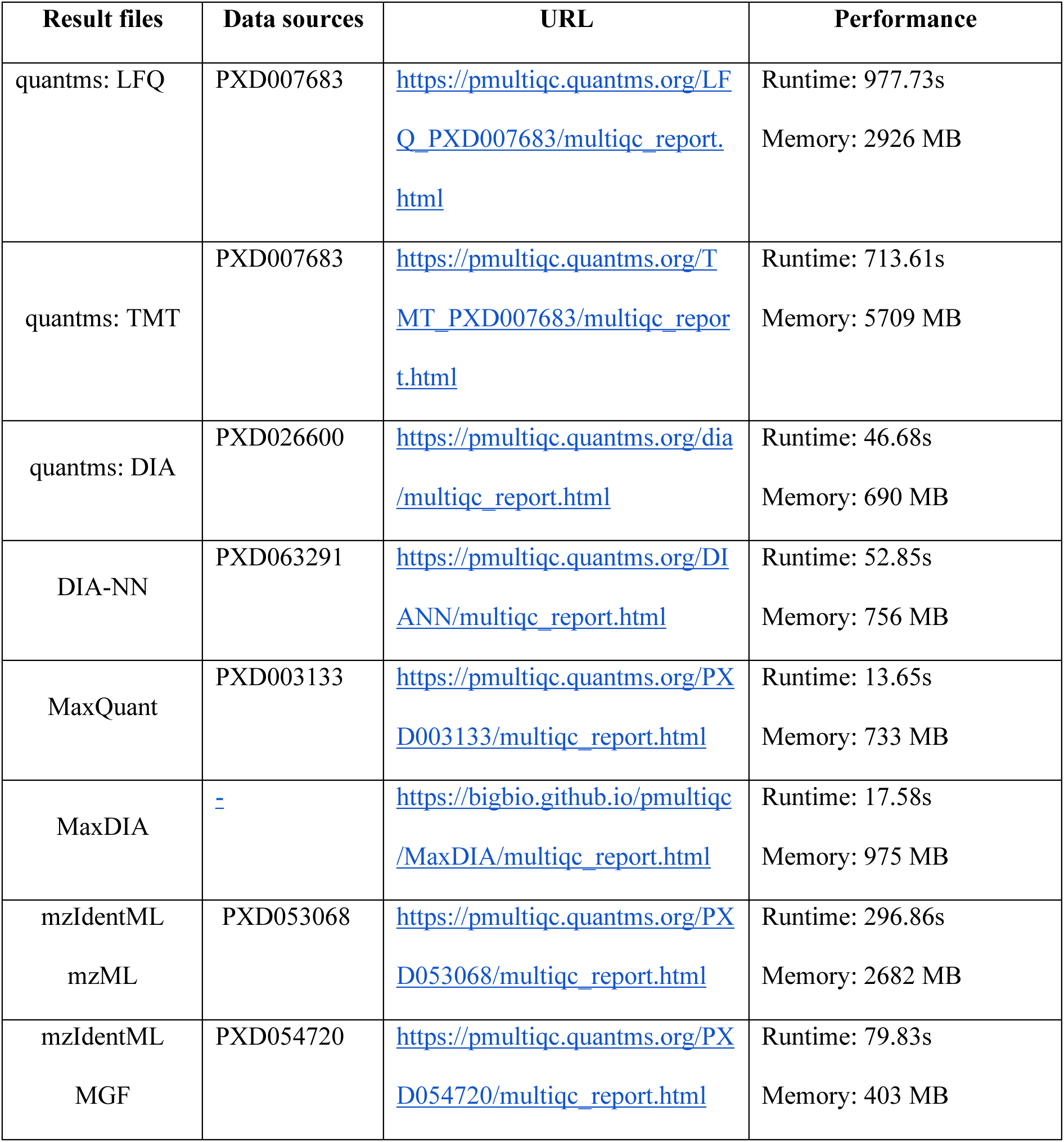
Example datasets for pmultiqc (https://pmultiqc.quantms.org/#-example-reports). The table lists the analysis tool used, a corresponding public dataset from PRIDE, and the computational resources and time required on a personal computer (CPU: Intel Core i9-7900X; OS: Ubuntu 24.04.2 LTS) to generate the QC report.

### Experimental design and metadata

The initial section of a pmultiqc report displays the experimental design and parameters in a table format. If sample metadata and experimental design are available for the dataset in SDRF (11) or any derived format (e.g., quantms *experimental_design.tsv*, https://pmultiqc.quantms.org/LFQ_PXD007683/multiqc_report.html#experiment_setup), a table showing the experimental design is presented, including samples and their relation to raw files. In addition, for MaxQuant result files, analysis parameters (recorded in *parameters.txt*) are directly converted into a table.

### Results overview

The first subsection, the *Summary Table*, summarizes the total number of acquired and identified MS2 spectra, the MS2 identification rate, the number of identified peptides, and the number of identified and quantified proteins. The *Summary Table* is available for all supported workflows and enables users to quickly review key figures of the analysis. A second high-level overview of the experiment is provided by the *QC Heatmap*, which displays the distribution of the following metrics for the experiment: *contaminants, peptide intensity, charge, missed cleavages, missed cleavages variability, identification rate over retention time, MS2 oversampling,* and *peptide missing values*. This heatmap has proven valuable for users of PTXQC, which quickly detects samples or raw files that exhibit atypical performance for specific metrics within an experiment (6, 20). pmultiqc summarizes the identification/quantification results according to experimental conditions (if SDRF is available) associated with different samples. The report includes statistics on the number of peptides, unambiguous peptides, modified peptides, and protein groups.

### Identification summary and search engine scores

The *Identification Summary* section, inspired by the PRIDE Inspector Toolsuite (5), presents critical metrics for evaluating the quality and reliability of peptide and protein identifications in bottom-up MS-based proteomics experiments. The number of peptides identified per protein reflects sequence coverage, which in turn impacts the confidence in protein identification and quantification reliability (21, 22). Proteins identified by multiple unique peptides (typically three or more) offer greater confidence, while single-peptide identifications (often called “one-hit wonders”) may arise by chance and require additional validation.

The missed cleavage distribution is a simple but effective measure of sample preparation quality, specifically reflecting how efficiently the enzymatic digestion process cleaves the proteins (23). An unusually high proportion of missed cleavages (>20% with one or more missed cleavages) may indicate incomplete digestion due to insufficient enzyme concentration, shortened digestion time, or the presence of digestion inhibitors. Well-prepared samples typically show >80% fully cleaved peptides, with systematic deviations suggesting the need for protocol optimization (1, 5, 24). The MS2 identification rate per raw file is another key quality metric; rates below 10% may indicate problems with sample preparation, database choice, or search settings, whereas rates above 40% generally reflect high-quality data. This section displays the search engine-specific scores for the peptide identification tool used, including posterior error probabilities (PEP) (**Figure 2A**), spectral E-values, cross-correlation scores (XCorr), and SAGE HyperScores (quantms multi search engine - https://pmultiqc.quantms.org/LFQ_PXD007683/multiqc_report.html#search_engine_scores).

**Figure 2:**
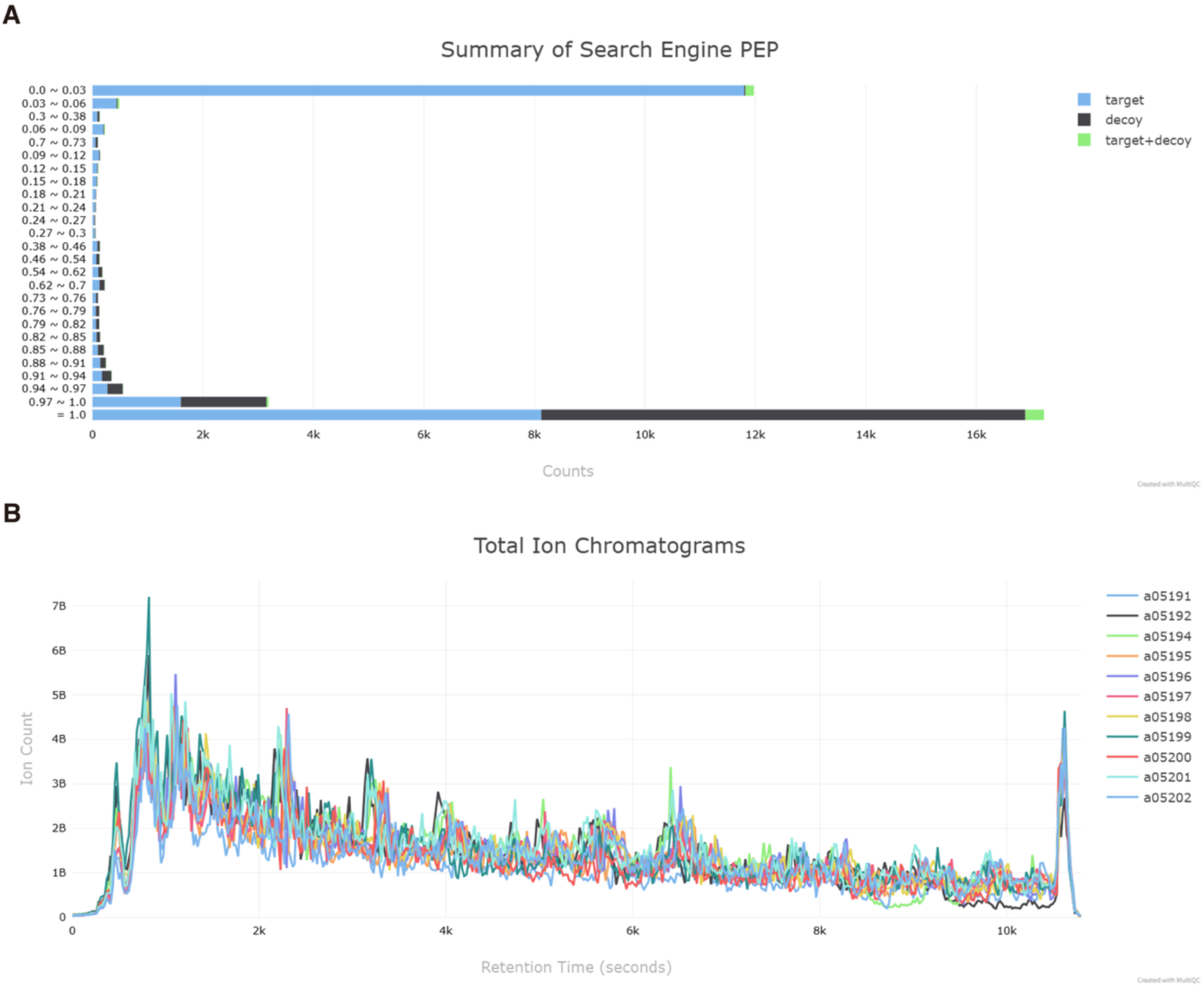
**(A)** Distribution of posterior error probabilities (PEP) from a search engine result. Bars represent the counts of peptide spectrum matchs (PSMs) classified as *target*, *decoy*, or *target+decoy* across different PEP intervals. The majority of identifications have low PEP values, reflecting high-confidence peptide assignments. (**B**) Total Ion Chromatograms (TICs) for all analyzed runs show consistent intensity profiles across MS runs, indicating uniform chromatographic performance and sample injection.

### Contaminants

Identification of common contaminants such as keratins, trypsin, and bovine serum albumin may indicate sample handling or reagent issues that may negatively affect data interpretation. Similar to PTXQC for MaxQuant, pmultiqc reports the top 5 contaminant proteins within the experiment. For each raw file (or group), all contaminants are identified, and their proportions are calculated based on intensity. In the quantms pipeline, contaminants are labeled by adding the prefix “CONTAMINANT_” to the accession. In MaxQuant, contaminants are indicated in the “Potential contaminant” field (entries marked with a “+” are considered contaminants).

### Quantification reports

pmultiqc provides visualizations to assess and interpret quantitative results including peptide intensity distributions, overall intensity distributions, label-free quantification (LFQ) intensity distributions, and principal component analysis (PCA) of both raw and LFQ intensities, which are log-transformed to ensure a more informative representation of the data (https://pmultiqc.quantms.org/PXD003133/multiqc_report.html#quantification_analysis). If the supported tool (e.g., MaxQuant) and analysis results contain peptide and protein quantification information, distributions of intensities, protein, and peptide quantitative results are summarized in tabular form.

### Retention time QC and mass errors

pmultiqc visualizes key features related to retention time in the subsection Retention time QC and mass errors. Specifically, it generates plots showing the overall distribution of retention times of every MS1 (DIA-NN, MaxQuant), or MS2 (quantms, mzIdentML) signals across all runs, as well as scatter plots depicting the relationship between retention time and peak width (i.e., the total retention time width of the peak), and between retention time and ion injection time. In addition, pmultiqc provides visualizations of mass error metrics, including delta mass (in (dalton) Da and parts per million (ppm)) and uncalibrated mass error (in ppm). For quantms pipelines, delta mass is calculated based on the difference between the experimental and theoretical mass-to-charge ratios (*m*/*z*), excluding uncalibrated mass error. In contrast, for MaxQuant pipelines, all of these mass error metrics are directly provided as part of the output. For the MaxQuant pipeline, we additionally followed the PTXQC approach to visualize TopN metrics, including their relationship with retention time.

### MS1 analysis

The MS1 analysis is primarily based on data extracted from spectrum file (*_ms_info.parquet) or in MaxQuant from the *msScans.txt*. pmultiqc processes this data to generate a series of analytical plots that provide a comprehensive overview of MS1 scan characteristics across all runs. First, total ion chromatograms (TIC) are produced for all analyzed runs, representing the total ion intensity at each time point during the MS1 scans. Total ion chromatograms (TICs) provide an overview of the ion accumulation profile across the chromatographic run. In contrast, MS1 base peak chromatograms (BPC) highlight the most intense ion signal at each time point, thereby emphasizing high-abundance compounds while minimizing baseline noise (**Figure 2B**). In addition, MS1 peak plots illustrate the number of peaks detected in each MS1 scan during a sample run, offering insights into spectral complexity and signal richness over time. Finally, a summary table titled “General Stats for MS1 Information” is generated, which includes acquisition date and time, total ion intensity for MS1, and total ion intensity for MS2 scans. This table provides a concise overview of the data quality and acquisition characteristics of each sample.

### MS2 and spectral statistics

pmultiqc also provides comprehensive statistical analyses of MS/MS spectra from both identified and unidentified scans. Specifically, three major aspects are examined: *number of peaks per MS/MS spectrum*, *peak intensity distribution*, and *distribution of precursor charges*. Each of these metrics is calculated separately for identified and unidentified spectra, allowing for comparative evaluation of spectral complexity, signal intensity, and charge state distribution across the dataset. It is important to note that for DIA data, only identified spectra are available and included in this analysis. Furthermore, pmultiqc generates a summary table titled “Pipeline Spectrum Tracking”, which tracks the number of MS1 and MS2 spectra, as well as the number of spectra identified by different search engines, including MS-GF+, Comet, and Sage. The table also reports the number of reliable peptide spectrum matches (PSMs) used for quantification and the number of quantified peptides that passed the final false discovery rate (FDR) thresholds at both the peptide and protein levels. In addition, pmultiqc performs per-raw-file statistics, summarizing the distribution of precursor charge states and the number of 3D peaks (defined as peaks corresponding to the same peptide sequence and the same charge state).

### Software versions and parameters

A key element of reproducible proteomics and adherence to FAIR principles (25) is the thorough documentation of software versions and analysis parameters. pmultiqc addresses this need by automatically extracting and presenting detailed version information and pipeline configurations from supported workflows. In the case of quantms, it generates comprehensive documentation that includes software versions, workflow parameters, and a standardized methods summary. The version tracking captures all tools used, such as MS-GF+, Comet, DIA-NN, and MaxQuant, ensuring reproducibility by accounting for even minor differences that can impact results. Pipeline parameters, including search settings, FDR thresholds, modifications, and quantification methods, are extracted directly from YAML files and mzTab metadata, ensuring consistency between documentation and actual analysis. Additionally, pmultiqc provides a citation-ready methods description that can be included directly in manuscripts, reducing the burden on researchers and promoting standardized reporting. For MaxQuant workflows, configuration details are extracted from the parameters.txt file, documenting key settings like search databases, enzyme specificity, and mass tolerances, thereby supporting both reproducibility and method comparison.

### pmultiqc Online Service

An additional service called *pmultiqc service* was developed to enable multiple groups and laboratories to run pmultiqc as a scalable, distributed service. The pmultiqc service leverages Redis (https://redis.io/) to manage jobs across distributed architectures with multiple nodes and supports flexible deployment options, including Kubernetes (k8s) and Docker Compose, enabling seamless integration into scalable computing environments. In addition to allowing deployment within your own company or research lab, we have deployed several distributed instances of pmultiqc, one of which is maintained by the PRIDE database team (https://www.ebi.ac.uk/pride/services/pmultiqc/), one by the Bioinformatics Solution Center at FU Berlin (https://pmultiqc.bsc.fu-berlin.de) and another by Tübingen University (https://abi-services.cs.uni-tuebingen.de/pmultiqc/). These instances enable users of pmultiqc to validate their data before submission to PRIDE or simply analyze their results. Users can upload a ZIP file containing result files from supported tools such as MaxQuant, DIA-NN, quantms, or mzIdentML plus peak list files.

Beyond generating reports from local data, the pmultiqc service also allows users to retrieve and analyze results directly from PRIDE datasets by providing a ProteomeXchange accession number. For example, users can submit an accession like PXD003133 to the service (https://www.ebi.ac.uk/pride/services/pmultiqc/submit?accession=PXD003133). This functionality helps public proteomics data users evaluate datasets before downloading or reusing them, even without extensive bioinformatics expertise. When a pmultiqc report is generated from local data or a PRIDE dataset, the pmultiqc service allows not only to download the results but also to inspect the generated reports in a browser.

### ProteoBench integration

In addition to supporting protein quantification pipelines, pmultiqc also enables the visualization of ProteoBench outputs. ProteoBench (https://proteobench.cubimed.rub.de/) is an open platform designed for benchmarking proteomics data analysis workflows, allowing for standardized performance evaluation across different tools and configurations. ProteoBench extracts from different protein quantification pipelines information on the peptide level into an intermediate data format, which is then passed by pmultiqc to generate reports. The feature is available for the quantification benchmarking modules, allowing a precomputed MultiQC report to be downloaded and visualized in the user’s browser (e.g. https://pmultiqc.quantms.org/ProteoBench/multiqc_report.html). The report contains sections that are more specific to the experimental setup in ProteoBench, following the idea of easy report customization. The report lists available information across conditions (species), which includes the number of identified precursor ions, intensity distributions, missing quantifications and identifications, coefficient of variations (CV), and log2 fold changes. These plots allow easy identification of rather low-quality submissions.

## Discussion

pmultiqc represents a major advancement in proteomics quality control by offering a unified, metadata-aware framework for comprehensive QC assessment across diverse analytical workflows. Building on the visualization capabilities of MultiQC and re-implementing metrics from tools like the PRIDE Inspector Toolsuite and PTXQC, pmultiqc introduces a lightweight, scalable solution for generating shareable web-based QC reports without requiring complex infrastructure. Its support for four major proteomics platforms, DIA-NN, MaxQuant, quantms, and PRIDE Complete submissions, demonstrates the feasibility of standardized QC reporting in a field known for its methodological diversity while preserving analytical depth and interpretability.

By combining comprehensive metric analysis with SDRF-driven metadata, pmultiqc enables more sophisticated quality assessment strategies that provide actionable insights for experimental optimization and troubleshooting. Its emphasis on reproducibility and FAIR compliance directly addresses key challenges in modern proteomics. Moreover, its flexible deployment options, including manual execution in Python, Docker containers, Docker Compose, and Kubernetes configurations, make it accessible for both individual labs and larger service facilities. Notably, the integration with public data repositories like PRIDE (https://www.ebi.ac.uk/pride/services/pmultiqc/) enables researchers to generate and share QC reports from multiple proteomics tools (e.g., DIA-NN, MaxQuant), facilitating data reanalysis and comparative quality assessment across studies.

pmultiqc’s modular design allows anyone to expand it by adding new workflows, tools, plots, QC metrics, or supported file formats. Extensive documentation and examples further lower the barrier for adoption and contribution. Looking ahead, pmultiqc’s metadata-aware approach should be extended as more tools support SDRF and more results are generated at the sample level, enabling the automatic generation of metrics without the need to combine run-based results into sample level results. Integration with benchmarking initiatives like ProteoBench could also enhance its role in performance evaluation and standardization. More importantly, the Quality Control Working Group of the HUPO-PSI developed the mzQC format (26) as a standard file format for exchanging, transmitting, and archiving quality metrics derived from MS. mzQC is growing its support by the community but a generic visualization of these metrics and the format is still missing. pmultiqc will help the community to fill that gap by providing an extendable, open-source library to support mzQC visualization reports. Ultimately, pmultiqc aims to boost and simplify generating and sharing QC reports in proteomics for popular tools, communities, and ProteomeXchange partners. By offering a consistent and accessible QC framework, it helps set baseline expectations for data quality across various experimental designs and analytical platforms.

## Data and Code Availability

The code for pmultiqc is available on GitHub as open source under the MIT license: https://github.com/bigbio/pmultiqc. In addition, the example QC reports can be read on the pmultiqc web page (https://pmultiqc.quantms.org), and all the example datasets can be found on the PRIDE database FTP server.

## Acknowledgments

T.S. and T.D.M acknowledges funding by the Federal Ministry of Education and Research in the frame of de.NBI/ELIXIR-DE (W-de.NBI-022). T.S. acknowledges support by the Ministry of Science, Research and Arts Baden-Württemberg (LIBIS). V.D. is supported by the German Ministry of Education and Research (BMBF), as part of the National Research Node “Mass spectrometry in Systems Medicine” (MSCoreSys), under grant agreement 161L0221. Y.P-R, S.K, C.B would like to acknowledge funding from EMBL core funding, Wellcome grants (208391/Z/17/Z, 223745/Z/21/Z) and the BBSRC grant ‘DIA-Exchange’ [BB/X001911/1].

## Author contributions

Q-X.Y., C.D., Y. P-R., T.S., C.B., S.K., C.B., developed the library and the service. T.D.M. deployed the service of pmultiqc in the Tubingen University. H.W., implemented the integration between pmultiqc and ProteoBench. A.L, W.B, J.U., J.X., P.E., J.C., V.D., O.K., M.B. tested the library reports and provided feedback about the metrics and plots that should be included for each independent tool. Y.P-R, M.B., T.S. wrote the main manuscript. All authors contributed to the manuscript, the design of the library, and discussions about the future of the open-source project.

## Abbreviations

BPC: Base Peak Chromatogram
CV: Coefficient of Variation
DDA: Data-Dependent Acquisition
DIA: Data-Independent Acquisition
Da: Dalton
FDR: False Discovery Rate
LFQ: Label-Free Quantification
MS: Mass Spectrometry
MS1: Mass Spectrometry level 1
MS2: Mass Spectrometry level 2
PEP: Posterior Error Probability
PCA: Principal Component Analysis
PTM: Post-Translational Modification
PSM: Peptide Spectrum Match
QC: Quality Control
TIC: Total Ion Chromatogram
m/z: Mass-to-Charge Ratio
ppm: Parts Per Million
SDRF: Sample and Data Relationship Format
XCorr: Cross-Correlation Score

## References

1. Neely, B. A., Perez-Riverol, Y., and Palmblad, M. (2024) Quality Control in the Mass Spectrometry Proteomics Core: A Practical Primer. J Biomol Tech 35

2. Bittremieux, W., Tabb, D. L., Impens, F., Staes, A., Timmerman, E., Martens, L., and Laukens, K. (2018) Quality control in mass spectrometry-based proteomics. Mass Spectrom Rev 37, 697–711

3. Perez-Riverol, Y., Bandla, C., Kundu, D. J., Kamatchinathan, S., Bai, J., Hewapathirana, S., John, N. S., Prakash, A., Walzer, M., Wang, S., and Vizcaino, J. A. (2025) The PRIDE database at 20 years: 2025 update. Nucleic Acids Res 53, D543–D553

4. Bittremieux, W., Valkenborg, D., Martens, L., and Laukens, K. (2017) Computational quality control tools for mass spectrometry proteomics. Proteomics 17

5. Perez-Riverol, Y., Xu, Q. W., Wang, R., Uszkoreit, J., Griss, J., Sanchez, A., Reisinger, F., Csordas, A., Ternent, T., Del-Toro, N., Dianes, J. A., Eisenacher, M., Hermjakob, H., and Vizcaino, J. A. (2016) PRIDE Inspector Toolsuite: Moving Toward a Universal Visualization Tool for Proteomics Data Standard Formats and Quality Assessment of ProteomeXchange Datasets. Mol Cell Proteomics 15, 305–317

6. Bielow, C., Mastrobuoni, G., and Kempa, S. (2016) Proteomics Quality Control: Quality Control Software for MaxQuant Results. J Proteome Res 15, 777–787

7. Dogu, E., Mohammad-Taheri, S., Abbatiello, S. E., Bereman, M. S., MacLean, B., Schilling, B., and Vitek, O. (2017) MSstatsQC: Longitudinal System Suitability Monitoring and Quality Control for Targeted Proteomic Experiments. Mol Cell Proteomics 16, 1335–1347

8. Trachsel, C., Panse, C., Kockmann, T., Wolski, W. E., Grossmann, J., and Schlapbach, R. (2018) rawDiag: An R Package Supporting Rational LC-MS Method Optimization for Bottom-up Proteomics. J Proteome Res 17, 2908–2914

9. Demichev, V., Messner, C. B., Vernardis, S. I., Lilley, K. S., and Ralser, M. (2020) DIA-NN: neural networks and interference correction enable deep proteome coverage in high throughput. Nat Methods 17, 41–44

10. Sinitcyn, P., Hamzeiy, H., Salinas Soto, F., Itzhak, D., McCarthy, F., Wichmann, C., Steger, M., Ohmayer, U., Distler, U., Kaspar-Schoenefeld, S., Prianichnikov, N., Yilmaz, S., Rudolph, J. D., Tenzer, S., Perez-Riverol, Y., Nagaraj, N., Humphrey, S. J., and Cox, J. (2021) MaxDIA enables library-based and library-free data-independent acquisition proteomics. Nat Biotechnol 39, 1563–1573

11. Dai, C., Fullgrabe, A., Pfeuffer, J., Solovyeva, E. M., Deng, J., Moreno, P., Kamatchinathan, S., Kundu, D. J., George, N., Fexova, S., Gruning, B., Foll, M. C., Griss, J., Vaudel, M., Audain, E., Locard-Paulet, M., Turewicz, M., Eisenacher, M., Uszkoreit, J., Van Den Bossche, T., Schwammle, V., Webel, H., Schulze, S., Bouyssie, D., Jayaram, S., Duggineni, V. K., Samaras, P., Wilhelm, M., Choi, M., Wang, M., Kohlbacher, O., Brazma, A., Papatheodorou, I., Bandeira, N., Deutsch, E. W., Vizcaino, J. A., Bai, M., Sachsenberg, T., Levitsky, L. I., and Perez-Riverol, Y. (2021) A proteomics sample metadata representation for multiomics integration and big data analysis. Nat Commun 12, 5854

12. Ewels, P., Magnusson, M., Lundin, S., and Kaller, M. (2016) MultiQC: summarize analysis results for multiple tools and samples in a single report. Bioinformatics 32, 3047–3048

13. Cox, J., and Mann, M. (2008) MaxQuant enables high peptide identification rates, individualized p.p.b.-range mass accuracies and proteome-wide protein quantification. Nat Biotechnol 26, 1367–1372

14. Dai, C., Pfeuffer, J., Wang, H., Zheng, P., Kall, L., Sachsenberg, T., Demichev, V., Bai, M., Kohlbacher, O., and Perez-Riverol, Y. (2024) quantms: a cloud-based pipeline for quantitative proteomics enables the reanalysis of public proteomics data. Nat Methods 21, 1603–1607

15. Gruning, B., Dale, R., Sjodin, A., Chapman, B. A., Rowe, J., Tomkins-Tinch, C. H., Valieris, R., Koster, J., and Bioconda, T. (2018) Bioconda: sustainable and comprehensive software distribution for the life sciences. Nat Methods 15, 475–476

16. da Veiga Leprevost, F., Gruning, B. A., Alves Aflitos, S., Rost, H. L., Uszkoreit, J., Barsnes, H., Vaudel, M., Moreno, P., Gatto, L., Weber, J., Bai, M., Jimenez, R. C., Sachsenberg, T., Pfeuffer, J., Vera Alvarez, R., Griss, J., Nesvizhskii, A. I., and Perez-Riverol, Y. (2017) BioContainers: an open-source and community-driven framework for software standardization. Bioinformatics 33, 2580–2582

17. Pfeuffer, J., Bielow, C., Wein, S., Jeong, K., Netz, E., Walter, A., Alka, O., Nilse, L., Colaianni, P. D., McCloskey, D., Kim, J., Rosenberger, G., Bichmann, L., Walzer, M., Veit, J., Boudaud, B., Bernt, M., Patikas, N., Pilz, M., Startek, M. P., Kutuzova, S., Heumos, L., Charkow, J., Sing, J. C., Feroz, A., Siraj, A., Weisser, H., Dijkstra, T. M. H., Perez-Riverol, Y., Rost, H., Kohlbacher, O., and Sachsenberg, T. (2024) OpenMS 3 enables reproducible analysis of large-scale mass spectrometry data. Nat Methods 21, 365–367

18. Levitsky, L. I., Klein, J. A., Ivanov, M. V., and Gorshkov, M. V. (2019) Pyteomics 4.0: Five Years of Development of a Python Proteomics Framework. J Proteome Res 18, 709–714

19. Harris, C. R., Millman, K. J., van der Walt, S. J., Gommers, R., Virtanen, P., Cournapeau, D., Wieser, E., Taylor, J., Berg, S., Smith, N. J., Kern, R., Picus, M., Hoyer, S., van Kerkwijk, M. H., Brett, M., Haldane, A., Del Rio, J. F., Wiebe, M., Peterson, P., Gerard-Marchant, P., Sheppard, K., Reddy, T., Weckesser, W., Abbasi, H., Gohlke, C., and Oliphant, T. E. (2020) Array programming with NumPy. Nature 585, 357–362

20. Schott, A. S., Behr, J., Geissler, A. J., Kuster, B., Hahne, H., and Vogel, R. F. (2017) Quantitative Proteomics for the Comprehensive Analysis of Stress Responses of Lactobacillus paracasei subsp. paracasei F19. J Proteome Res 16, 3816–3829

21. Muntel, J., Boswell, S. A., Tang, S., Ahmed, S., Wapinski, I., Foley, G., Steen, H., and Springer, M. (2015) Abundance-based classifier for the prediction of mass spectrometric peptide detectability upon enrichment (PPA). Mol Cell Proteomics 14, 430–440

22. Audain, E., Uszkoreit, J., Sachsenberg, T., Pfeuffer, J., Liang, X., Hermjakob, H., Sanchez, A., Eisenacher, M., Reinert, K., Tabb, D. L., Kohlbacher, O., and Perez-Riverol, Y. (2017) In-depth analysis of protein inference algorithms using multiple search engines and well-defined metrics. J Proteomics 150, 170–182

23. Lawless, C., and Hubbard, S. J. (2012) Prediction of missed proteolytic cleavages for the selection of surrogate peptides for quantitative proteomics. OMICS 16, 449–456

24. van den Broek, I., Smit, N. P., Romijn, F. P., van der Laarse, A., Deelder, A. M., van der Burgt, Y. E., and Cobbaert, C. M. (2013) Evaluation of interspecimen trypsin digestion efficiency prior to multiple reaction monitoring-based absolute protein quantification with native protein calibrators. J Proteome Res 12, 5760–5774

25. Wilkinson, M. D., Dumontier, M., Aalbersberg, I. J., Appleton, G., Axton, M., Baak, A., Blomberg, N., Boiten, J. W., da Silva Santos, L. B., Bourne, P. E., Bouwman, J., Brookes, A. J., Clark, T., Crosas, M., Dillo, I., Dumon, O., Edmunds, S., Evelo, C. T., Finkers, R., Gonzalez-Beltran, A., Gray, A. J., Groth, P., Goble, C., Grethe, J. S., Heringa, J., t Hoen, P. A., Hooft, R., Kuhn, T., Kok, R., Kok, J., Lusher, S. J., Martone, M. E., Mons, A., Packer, A. L., Persson, B., Rocca-Serra, P., Roos, M., van Schaik, R., Sansone, S. A., Schultes, E., Sengstag, T., Slater, T., Strawn, G., Swertz, M. A., Thompson, M., van der Lei, J., van Mulligen, E., Velterop, J., Waagmeester, A., Wittenburg, P., Wolstencroft, K., Zhao, J., and Mons, B. (2016) The FAIR Guiding Principles for scientific data management and stewardship. Sci Data 3, 160018

26. Bielow, C., Hoffmann, N., Jimenez-Morales, D., Van Den Bossche, T., Vizcaino, J. A., Tabb, D. L., Bittremieux, W., and Walzer, M. (2024) Communicating Mass Spectrometry Quality Information in mzQC with Python, R, and Java. J Am Soc Mass Spectrom 35, 1875–1882

