## Supplementary Notes for "pmultiqc: An open-source, lightweight, and metadata-oriented QC reporting library for MS proteomics"

^5^ Department of Biochemistry and Molecular Biology. Faculty of Science and Technology. University of the Basque Country (UPV/EHU), Bilbao, Spain

^6^ Department of Computer Science, University of Antwerp, 2020 Antwerp, Belgium.

^7^ Ruhr University Bochum, Medical Faculty, Medical Bioinformatics, 44801 Bochum, Germany

^8^ Ruhr University Bochum, Medical Faculty, Core Unit Bioinformatics - CUBiMed.RUB, 44801 Bochum, Germany.

^9^ Department of Computer Science, Applied Bioinformatics, University of Tübingen, Tübingen, Germany.

^10^ Computational Systems Biochemistry, Max Planck Institute of Biochemistry, 82152 Martinsried, Germany.

^11^ Seqera, Carrer de Marià Aguiló, 28, Barcelona, 08005, Spain.

^12^ Quantitative Proteomics laboratory, Charité – Universitätsmedizin Berlin, Berlin, Germany

^13^ Institute for Bioinformatics and Medical Informatics, University of Tübingen, 72076 Tübingen, Germany.

^14^ Institute for Translational Bioinformatics, University Hospital Tübingen, 72074 Tübingen, Germany.

^15^ Bioinformatics Solution Center, Institute of Computer Science, Freie Universität Berlin, 14195 Berlin, Germany.

### **Supplementary Note 1: List of metrics and sections implemented in pmultiqc.**

**Table 1**: List of QC metrics implemented in pmultiqc, with the given section, description of the metric, the workflow that supports the given metric, and some relevant citations of studies and tools where the metric has been previously used. QC metrics in pmultiqc reports are organized into sections, helping the users of the tool to understand the flow of the experiment.

| **Metric Name** | **Section** | **Description** | **Supported Workflows** | **Citation** |
| --- | --- | --- | --- | --- |
| Experiment Design | Experimental Design and Metadata | Shows experimental conditions, biological replicates, fractions, and sample metadata | quantms,  DIA-NN, MaxQuant | PMID: 28802010 |
| MaxQuant Parameters | Experimental Design and Metadata | Complete MaxQuant analysis parameters and settings | MaxQuant | PMID:26653327 |
| Summary Statistics | Results Overview | Overall pipeline statistics, including total MS2 spectra, identification rates, and peptide/protein counts | quantms, MaxQuant, DIA-NN | PMID:28802010 |
| Peptide ID Count | Identification Summary | Distribution of peptides per protein group | quantms, MaxQuant, DIA-NN | PMID:26653327 |
| Protein Groups Count | Identification Summary | Statistics on protein group identifications | quantms, MaxQuant, DIA-NN | PMID:26653327, PMID: 26545397 |
| Missed Cleavages | Identification Summary | Analysis of enzymatic digestion efficiency by file | quantms, MaxQuant | PMID:24760958, PMID: 26545397 |
| Modifications | Identification Summary | Distribution of post-translational modifications | quantms, MaxQuant, DIA-NN | PMID:28802010, PMID: 26545397 |
| MS/MS Identified | Identification Summary | Identification success rates per raw file | quantms, MaxQuant | PMID:26974716, PMID: 26545397 |
| Search Engine Scores | Identification Summary | Distribution of search engine scoring metrics (E-values, cross-correlation, PEP) | quantms | PMID:24760958 |
| Consensus Across Engines | Identification Summary | Agreement between multiple search engines | quantms | PMID:28802010 |
| Peptide Quantification | Quantification Analysis | Quantitative intensities for identified peptides across conditions | quantms, MaxQuant, DIA-NN |  |
| Protein Quantification | Quantification Analysis | Protein-level quantitative data across experimental conditions | quantms, MaxQuant, DIA-NN |  |
| Peptide Intensity Distribution | Quantification Analysis | Statistical distribution of peptide intensities | quantms, MaxQuant, DIA-NN | PMID:26974716 |
| Intensity Standard Deviation | Quantification Analysis | Variability analysis of quantitative measurements | DIA-NN |  |
| Total Ion Chromatograms | MS1 Analysis | Overall ion current across the acquisition time | quantms, DIA-NN | PMID:24760958 |
| Base Peak Chromatograms | MS1 Analysis | The most intense peak across the retention time | quantms, DIA-NN | PMID:24760958 |
| MS1 Statistics | MS1 Analysis | Summary statistics for MS1 data, including acquisition dates and current | quantms, DIA-NN | PMID:28802010 |
| MS2 Peak Distribution | MS2 and Spectral Stats | Distribution of peaks per MS/MS spectrum | quantms | PMID:26974716, PMID: 26545397 |
| Peak Intensity Distribution | MS2 and Spectral Stats | Statistical distribution of peak intensities | quantms,  DIA-NN | PMID:26974716 |
| Pipeline Spectrum Tracking | MS2 and Spectral Stats | Detailed tracking of spectra through pipeline stages | quantms | PMID: 26545397 |
| Precursor Charge Distribution | MS2 and Spectral Stats | Distribution of precursor ion charges by file | quantms,  DIA-NN | PMID:24760958, PMID: 26545397 |
| MS/MS 3D-peak Counts | MS2 and Spectral Stats | Analysis of MS/MS counts per detected feature | quantms,  MaxQuant | PMID:28802010 |
| IDs over RT | RT Quality Control | Identification success across retention time | quantms,  DIA-NN,  MaxQuant | PMID:28802010 |
| Delta Mass Analysis | Mass Error Trends | Mass measurement accuracy in Da and ppm | quantms,  DIA_NN,  MaxQuant | PMID:24760958, PMID: 26545397 |
| Software Versions | Software Versions | Complete listing of software tools and versions used | quantms | PMID:27312411 |
| Workflow Summary | Workflow Summary | Complete pipeline parameters and configuration | quantms | PMID:27312411 |
| Methods Description | Methods | Standardize the methods section text with proper citations | quantms | PMID:27312411 |

### **Supplementary Note 2: metrics and plots**

**Table 2: Statistics for the plot section**

| Sections | Sub sections | Plot types |
| --- | --- | --- |
| Experimental Design and Metadata | Experimental Design ^a, b, c, d^ | table |
|  | Parameters ^d^ | table |
| Results Overview | Summary Table ^a-e^ | table |
|  | HeatMap ^a-e^ | heatmap |
|  | Pipeline Result Statistics ^a-c, e^ | table |
| Identification Summary | Number of Peptides identified Per Protein ^a-e^ | bar |
|  | ProteinGroups Count ^a-e^ | bar |
|  | Peptide ID Count ^a-e^ | bar |
|  | Missed Cleavages Per Raw File ^a, d^ | bar |
|  | Modifications Per Raw File ^a-d^ | bar |
|  | MS/MS Identified Per Raw File ^a, d, e^ | bar |
| Search Engine Scores | Summary of Spectral E-values ^a^ | bar |
|  | Summary of cross-correlation scores ^a^ | bar |
|  | Summary of Search Engine PEP ^a^ | bar |
|  | Summary of Hyperscore ^a^ | bar |
|  | Consensus Across Search Engines ^a^ | bar |
|  | Summary of Andromeda Scores ^d^ | bar |
| Contaminants | Top5 Contaminants Per Raw File ^a, d^ | bar |
|  | Potential Contaminants Per File ^a, d^ | bar |
| Quantification Analysis | Peptides Quantification Table ^a-e^ | table |
|  | Protein Quantification Table ^a-e^ | table |
|  | Intensity Distribution ^a-d^ | box graph |
|  | PCA of Intensity ^d^ | scatter |
| MS1 Analysis | Total Ion Chromatograms ^a-c, e^ | line graph |
|  | MS1 Base Peak Chromatograms ^a-c, e^ | line graph |
|  | MS1 Peaks ^a-c, e^ | line graph |
|  | General stats for MS1 information ^a-c, e^ | table |
| MS2 and Spectral Stats | Number of Peaks per MS/MS spectrum ^a-c, e^ | bar |
|  | Peak Intensity Distribution ^a-c, e^ | bar |
|  | Pipeline Spectrum Tracking ^a^ | table |
|  | Distribution of Precursor Charges ^a-c, e^ | bar |
|  | Charge-state of Per File ^a-e^ | bar |
|  | MS/MS Counts Per 3D-peak ^a, d, e^ | bar |
| Mass Error Trends | Delta Mass [Da] ^a-d^ | line graph |
|  | Delta Mass [ppm] ^a, d^ | line graph |
|  | Uncalibrated Mass Error ^d^ | box graph |
| RT Quality Control | IDs over RT ^a-e^ | line graph |
|  | Normalisation Factor over RT ^b, c^ | line graph |
|  | FWHM over RT ^b, c^ | line graph |
|  | Peak width over RT ^b-d^ | line graph |
|  | Absolute RT Error over RT ^b, c^ | line graph |
|  | LOESS RT ~ iRT ^b, c^ | line graph |
|  | Ion Injection Time over RT ^d^ | line graph |
|  | TopN over RT ^d^ | line graph |
|  | TopN ^d^ | bar |
| quantms Software Versions | -^a, b^ | - |
| quantms Workflow Summary | -^a, b^ | - |
| quantms Methods Description | -^a, b^ | - |

a: quantms LFQ and TMT; b: quantms DIA; c: DIA-NN; d: MaxQuant (MaxDIA); e: mzIdentML
